# maniFasta and the AllOralsDB: simplifying construction of comprehensive reference databases for metaproteomics

**DOI:** 10.64898/2026.08.08.739415

**Authors:** Christopher Handelmann, Ashley K. Miles, Yinyin Ye, Marcelo Freire, Floyd E. Dewhirst, Tsute Chen, Jessica Mark Welch, Kathryn M. Kauffman

## Abstract

Metaproteomics aims to capture a taxonomically comprehensive snapshot of proteins in a sample. Design of reference databases is a key aspect of metaproteomic workflows, as these define what is ultimately seen. Databases tailored to focal biomes offer optimal performance, yet their construction often requires drawing on heterogeneous data sources, posing a challenge to reproducibility and documentation. Here we present maniFasta, a tool enabling users to generate standardized, reproducible, and robustly documented protein reference sets from diverse input sources and datatypes. Users provide information on their desired input types and sources, and the output is an integrated database comprising a protein sequence file (FASTA), with harmonized identifiers and standardized headers, and an associated provenance metadata table (manifest). We highlight the value of maniFasta in the context of salivary metaproteomics, addressing the need for a taxonomically comprehensive reference database. The AllOralsDB resource includes human proteins, as well as proteins from bacteria and archaea, fungi and other microeukaryotes, viruses and viroid-like elements, dietary sources, and common contaminants. Together, this work provides a community resource for oral and salivary metaproteomics (https://www.homd.org/ftp/AllOralsDB/), and a versatile and accessible tool for constructing protein databases for metaproteomics generally (https://github.com/KauffmanLab/maniFasta).

## INTRODUCTION

Metaproteomics can provide a snapshot of the landscape of proteins in a given sample at a given time^1,2^ - and holds potential to offer insight into interactions, and to mechanisms underpinning observed features of microbial community structure and function. Design of optimal reference databases, that enable accurate characterization of sample composition, is thus an integral aspect of metaproteomics^3,4^. In traditional database-search metaproteomics, reference sequences define the initial search space for identifying spectra; in *de novo* approaches, reference databases are used to assign predicted peptides to candidate source proteins and taxa. Database construction is therefore a critical step, and studies of database design have highlighted the value of sample- or biome-specificity in terms of ensuring accuracy of attributions, decreasing false discovery rates, and increasing computational efficiency^3–7^.

A recent review by Arikan and Atabay highlighted key challenges and practical recommendations for the construction of protein sequence databases for metaproteomics^3^. A major theme is that biome specificity is important, yet constructing databases that can strike the balance of being comprehensive without being overly large remains operationally challenging. Tools including ConDiGA^8^, gNOMO2^9^, MAPLE^10^, mPies^11^, ProteoClade^12^, Unipept Desktop 2.0^13,14^, metaProDB^15^, and MetaNovo^16^, and others have been developed to facilitate construction of databases, but are limited to specific input types. There is also an increasing need to collect reference proteins from various key public protein databases, including general repositories (e.g. NCBI, UniProtKB, RefSeq^17^, proGenomes3^18^, MGnify^19^) and biome-specific databases (e.g. *e*HOMD^20^, MOMD^21^, OM-RGC^22^, UHGP Catalogue^23^, and MetaPep^24^), and diverse others highlighted by Rigden and Fernández in the 2026 Nucleic Acids Research database issue^25^. While it is important to encompass the full range of relevant taxa, and integration of protein sequences from multiple sources is a useful approach for achieving this, a challenge is that a tool that facilitates integration and harmonization of heterogeneous source data for database construction is lacking^3^.

An additional consideration, beyond the impact of database construction on a given study, is the importance of reproducibility in enabling cross-study comparisons. This was highlighted in a recent perspective piece by Van Den Bossche et al.^26^, which underscored the need for standardization and improved open meta(data) practices in metaproteomics, including in database construction.

Here, we present maniFasta, a tool that accomplishes the deceptively “simple” task of creating reproducible, well-documented protein reference databases from diverse data types, including lists of species names, taxon identifiers, genome/proteome/protein accessions, and protein sequences. Public repositories currently supported by maniFasta include NCBI, UniProt Knowledgebase (UniProtKB), UniProt Archive (UniParc), Protein Data Bank (PDB), and the expanded Human Oral Microbiome Database (eHOMD)^20^. Design principles guiding the development of maniFasta include an open source community-evolvable code base, accessibility to users through both command line and web-interfaces, and an emphasis on robust documentation of input data and build parameters that follow FAIR Principles^27^ (Findable, Accessible, Interoperable and Reusable) for data management. Our overarching aim with this tool is to ease construction and documentation of reference protein databases and thereby to support taxonomically comprehensive metaproteomic analyses, as well as facilitate comparisons across studies.

We note that the design and development of maniFasta was inspired by challenges encountered during our initial efforts to generate a taxonomically comprehensive reference protein database for the field of salivary and oral metaproteomics. Thus, we present both maniFasta and our flagship use case, the AllOralsDB, which is intended to serve as an evolving resource for the salivary and oral metaproteomics community. Together, maniFasta and the AllOralsDB offer an example of the value of flexible, yet standardized, database construction for generating evolvable community metaproteomic reference sets.

## METHODS

Code for maniFasta is written in bash and python and was developed for use in a Unix environment. To ensure broad accessibility, a Google Colab notebook version of the tool is available, which allows generation of maniFasta databases using an interactive graphical user interface. The Colab notebook version guides the user through each step of the build design and construction, culminating with generation of a downloadable zip file of the user-designed database and all associated run logs and build documentation. In both cases, when used at command line or through the Colab GUI, the code is sourced from the associated GitHub repository. Further details are provided below with description of the tool.

Design of the AllOralsDB taxon composition was guided by review of the literature, and builds on the foundation of oral bacterial genomes defined in the *e*HOMD. As the *e*HOMD includes millions of proteins, we designed maniFasta to enable filtering to include subsets of sequences in a taxon-rank-dependent fashion.

To compare run times and performance across different flavors of the AllOralsDB we use FragPipe (https://github.com/Nesvilab/FragPipe). The selected sample is taken from the OSample study^28^, which optimized laboratory methods for detection of microbes in salivary metaproteomes. We select a sample produced using the method determined in their study to be optimal for recovery of microbial proteins [LUMOS2_20230602_H_Saliva_GX_S_R1_78min2.raw; ProteomeXchange ID: PXD055269, iProX ID: IPX0009371001]. Analyses with FragPipe (v24.0) were performed with the precursor mass tolerance set to -20 to 20 PPM and a fragment tolerance of 0.5 Da to fit the samples’ low-resolution ion-trap MS/MS. Peptides were required to be fully tryptic (strict trypsin, C-terminal to K/R) with up to two missed cleavages, 7–50 residues, and 500–5,000 Da. Carbamidomethylation of cysteine was specified as a fixed modification, and oxidation of methionine (+15.9949 Da) and protein N-terminal acetylation as variable modifications. Decoy reverse protein sequences were generated for each protein database utilizing the built-in FragPipe option; each database was split into 32 partitions and searches were run with 12 GB of RAM on 8 threads using a Windows 10 workstation with Intel Core i9-12900H with 20 logical cores. Protein inference and filtering were performed with ProteinProphet^29^ (v6.0.0-rc15) in Philosopher^30^ (v5.1.3-RC9), using razor-peptide assignment for shared peptides with a protein FDR set to 1% and using Philosopher’s 1% default for PSM, peptide and ion levels The AllOralsDB.v2026.215 build series (A, S, G, and F) was used for searches, and is preserved and available for download at *e*HOMD (see Data Availability). The runtimes for each flavor were supplied by FragPipe upon each run completion.

## RESULTS

### maniFasta - a flexible tool for construction of metaproteomic reference databases

The central purpose of the maniFasta toolkit is to enable users to easily construct comprehensive databases by integrating protein sequences from heterogeneous source types in a reproducible and robustly documented manner. maniFasta is unique in accepting a wide range of input types, including simple lists of species names, taxon identifiers, genome and protein accessions, as well as FASTA files, and also offers special handling of special data collections. In brief, users provide information about each of their desired input types, populate required fields in configuration files, and launch the pipeline, which then outputs the harmonized database FASTA file and metadata manifest, as well as packages documentation of run inputs, scripts, summary files, and logs (Figure 1).

**Figure 1.**
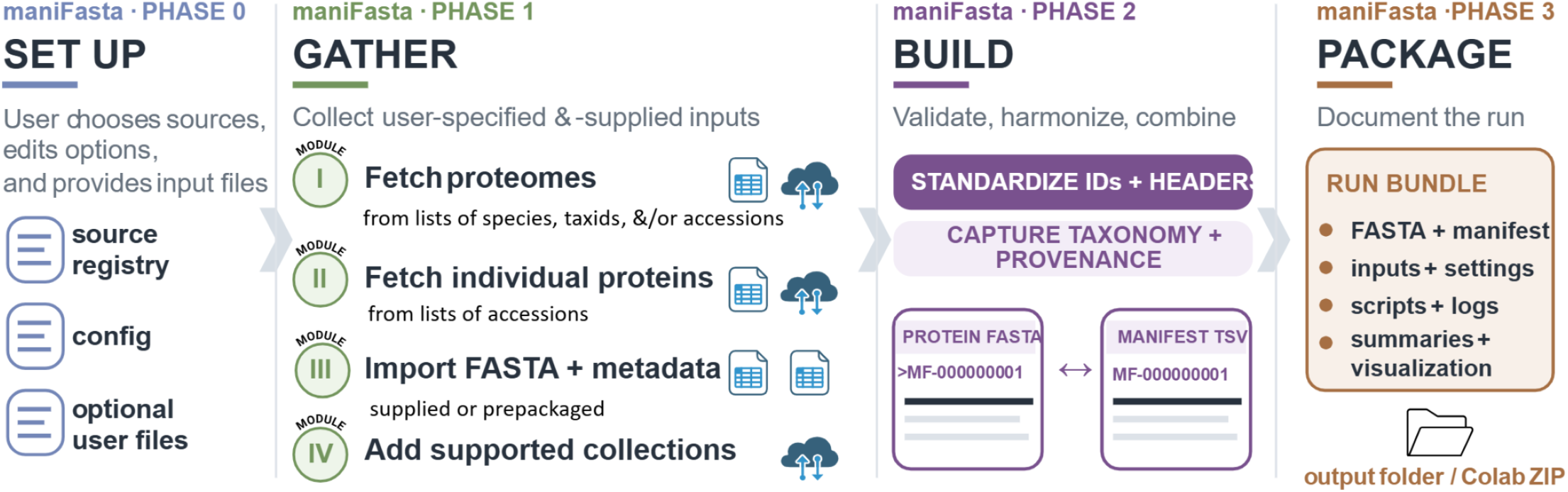
Overview of the maniFasta toolkit, highlighting the four phases spanning setup, collection of supplied and specified proteins, construction of core database files, and packaging of the run and associated documentation.

The tool can be run either at the command line or through an interactive web page, in both cases by sourcing from the maniFasta GitHub repository. Users comfortable with the command line can build their database locally on their machine or on a compute cluster. Users who prefer to generate a database directly online through an interactive interface, can do so by accessing the maniFasta Google Colab notebook website (see Data Availability, below). The Colab notebook guides the user in uploading files and editing the configuration files and then builds the database, generating all outputs for direct download.

Regardless of where it is launched, the maniFasta pipeline proceeds in four phases, as follows:

In **Phase 0** the user sets up the run by editing configuration files and providing the relevant input files to support various desired data type integrations. All setup occurs in the 00.setup directory, which contains three configuration files that define local tool paths (000.maniFasta.local.env), variables (000.maniFasta.config), and the input types requested or provided by the user (000.maniFasta.source_registry.tsv). The user then launches Phase 1 by either running runBuild_db.sh directly (if on a personal machine), by submitting 01.submit_runBuild_db.sh (if running on a compute cluster), or by clicking to run “Cell 1” (if using Google Colab).

In **Phase 1** the desired database components are automatically collected and staged by maniFasta, per variables defined by source_planner.py from the source_registry, and using one or more of four available Modules (described below). For all modules (Table 1), a key element of the maniFasta approach is the systematic capture of taxon and source annotation data for integration into the database manifest. The following modules are currently represented.

**Table 1.** maniFasta Phase 1 module inputs, processes, and outputs, and list of included pre-packaged inputs. Following Phase 0 setup of the indicated inputs, maniFasta gathers all sequences in Phase 1.

| MODULE | INPUTS | PROCESS | OUTPUT | PREPACKAGED INPUTS |
| --- | --- | --- | --- | --- |
| **Module I**<br>**proteome fetch**<br>(NCBI, UniProt) | TSV file(s) with columns for: species, taxid, accession, fetch_source, note. One of species/taxid/accession must be filled; fetch_source and note are optional. | For each row, a representative genome for the indicated species/taxid/accession is identified and all associated protein records are downloaded. Default fetch_source unless otherwise specified is NCBI, UniProtKB fetches also accept a UniProt proteome ID (UPID) and a proteome type: best (default), reference, representative, any, or pan. | Fetches proteome sequences with harmonized taxonomy and source metadata. | Curated accession lists assembled for the AllOralsDB (see Table 3). |
| **Module II**<br>**protein fetch**<br>(NCBI, UniProt, UniParc, PDB) | TSV files(s) with columns for: accession (required), fetch_source, note. May include mixed fetch_sources. | TSV file(s) with a list of individual protein accessions — a single mixed file or per-source; a per-row fetch_source sets the backend: ncbi (default), uniprot, uniparc, or pdb. | Fetches protein sequences with harmonized taxonomy and source metadata. | AllergenOnline v24 (accessions of 2,373 food- and environmental-associated allergens); HSP 2.0 salivary proteins (accessions of 3,437 proteins in the Human Salivary Proteome knowledgebase, downloaded June 18, 2026.). |
| **Module III**<br>**FASTA**<br>**sequences** | FASTA file plus an optional metadata TSV with columns for protein_id (required and must match FASTA), source, ncbi_taxid, genus, species, description. Without metadata the defaults are taxid 32644 (unknown) and genus/species NA/NA. | Protein sequences are integrated into the harmonized maniFastaDB. | Integrated sequences with user-supplied or pre-packaged metadata. | Dairy_DB (dataset of 171 mammalian milk protein sequences curated by Hendy 2019 https://doi.org/10.15124/589742eb-287a-4576-a00a-30df33d9f52c); *Trichomonas tenax* protein sequences (Mpeyako et al. 2024 Suppl. Table 2 & Suppl. Data 3); Obelisk protein sequences (11,581), encoded on Obelisk genomes provided in Zheludev et al. 2024 Supplementary File 1 |
| **Module IV**<br>**collections** | Selection of desired options in the source_registry TSV and config files. | Fetches per user-specifications for supported collections: Human [UniProt human proteome — canonical (default), canonical + isoforms, or + TrEMBL]; bacterial and archaeal: HOMD [release version; optional oral body-site filter; optional species / genus / family dereplication]; contaminants: cRAP (CCP). | Integrated collection proteins with associated metadata. | UniProtKB human proteome collections (https://www.uniprot.org/proteomes/UP000005640); HOMD bacterial and archaeal proteomes (https://hombd.org/); Cambridge Centre for Proteomics contaminant set (https://zenodo.org/records/15115102). |

**Module I** performs proteome-level data fetching from public repositories, including NCBI and/or UniProtKB, based on user preferences. Users can specify the desired proteome with an accession ID (highest specificity), then taxonomy identifier (TaxID; note UniProtKB OX=NCBI_TaxID), and/or a species name (lowest specificity). Supplying any of these will fetch the specified proteome, but including multiple fields allows the module to source the proteome in the case of an issue such as an outdated accession. When supplied with multiple pieces of information for a proteome source, the module proceeds from the most specific to the least (accession > TaxID > species name). The download schemas are similar for either the NCBI or the UniProtKB sources, with a few specific options based on the selected source, as described below. When using this module it is important to note that proteome accessions are a specific entity and thus module I will only work if UniProtKB is using a defined proteome.

We further allow for the user to specify a species pan proteome, sourcing proteins from multiple strains in a species to represent both the core proteins of a species along with variable and unique proteins found in only a subset of strains or individuals^31^. This curation is available from UniProtKB where the pan proteome IDs are based on the TaxID with pp appended to the front of it (i.e. the TaxID for *E. coli* is 562, the pan proteome is pp562). To retrieve one or more pan proteomes from UniProtKB, the user supplies the TaxIDs in a file that they add to the 00.setup directory and enter as a new row in the source_registry. If specified to source the pan proteome “fetch_source=uniprot;uniprot_proteome_type=pan” then the program will append pp onto the TaxID and use this to source the pan proteome. Module I also allows the user to specify a higher taxon TaxID, allowing the pipeline to select the first reference proteome for the taxon level that is retrieved to be sourced and downloaded as a representative for the taxon at that specified level.

For NCBI, if an assembly accession is provided, the system retrieves the corresponding TaxID, organism, and strain directly from the NCBI Datasets assembly summary and selects that assembly for download. If no accession is provided, a TaxID is resolved to an organism name using NCBI E-utilities (efetch), and a species name is mapped to a TaxID via an E-utilities search (esearch)^32^. In both instances, the organism is queued to download the best available assembly. If the assembly-accession lookup fails, the process falls back to the user-supplied TaxID, and then to the species name when available. After determining the download plan, the proteomes are fetched. Organisms resolved by TaxID or name are fetched using NCBI Datasets (ncbi-datasets-cli), selecting the single best available assembly according to the preference order: 1) RefSeq reference, 2) RefSeq representative, 3) any RefSeq, and then 4) GenBank, using the first that returns protein annotations. Organisms with a successful user-specified assembly accession bypass this selection and download the named assembly directly. The 000.maniFasta.local.env allows the user to set the NCBI email and API key.

For UniProtKB sources, the user can provide a specific UniProtKB ID (UPID) for download, along with a taxonomic identifier (TaxID) and/or species name. The user can further specify the proteome type (“best” by default, “reference”, “representative”, “any”, or “pan”). The best option searches through a ranked hierarchy within UniProtKB categories: 1) Reference, 2) Representative, 3) Other, and then 4) Redundant and returns the top-ranking proteome for any given species name or TaxID. UniProtKB also allows isoforms to be included (default is not to include isoforms) and lets the user select whether the UniProtKB selection is reviewed (default, both reviewed [Swiss-Prot] and unreviewed [TrEMBL] proteins are included; selecting true will return only reviewed proteins). Both paths download the proteomes and record each download in the proteomes_manifest.tsv file.

**Module II** is a protein-level fetch from NCBI, UniProtKB, PDB, and/or UniParc per user preference; users provide a list of individual protein accessions, this list is then fetched and integrated into the database. Users may opt to include pre-packaged accession lists for fetch, which currently include proteins identified in human saliva (described in detail below) and proteins identified as food- and environmental-associated allergens in the AllergenOnlineV24 database^33^.

The individual protein accessions can be provided by the user as a single file with multiple source types, or as sets of accessions grouped by source in separate files. The list of protein accessions from various sources is read per row, using the fetch_source for each row to determine the protein source, defaulting to NCBI. The NCBI protein accessions are fetched by the eUtils efetch^32^. The UniProtKB accession is sourced from “rest.uniprot.org/uniprotkb/accessions“ directly and the TaxID is pulled from the UniProtKB FASTA header. The UniParc source is accessed via “rest.uniprot.org/uniparc/{UPI}.json” but may lack a specific species, as a single UPI can map to multiple organisms^34^. The pipeline checks for the presence of a TaxID and specifies an origin down to the specific genus and species. If the protein cannot be attributed to a single species, then the TaxID and species are selected based on the taxon with the highest count in the pool. The pool can be modified by the user specifying a pick mode: “best”, “common”, or “first”. Selecting “best” narrows the species to active and then pools the taxa in the highest hierarchical order of: 1) Swiss-Prot, 2) TrEMBL/UniProtKB, and 3) RefSeq. Active sources are always preferred to obsolete ones, so an active source from RefSeq will be used over an obsolete taxon from Swiss-Prot. Selecting “common” removes the ranking so all taxon sources are pooled to determine the most common taxon, and “first” gives the first taxon fetched by the API. All UPI results return an “n_organisms=“ showing how many taxa were matched to the UPI, with more than one matched species generating the flag “ambiguous_organism”. If the UPI lacks a TaxID or full binomial then the TaxID is left blank and the genus and species are labeled NA/NA. Protein sequences are sourced from PDB per structure entry from RCSB FASTA using www.rcsb.org/fasta/entry/{ENTRY}^35^. The PDB accessions can be given either as entry-level (i.e. 5Y36), returning all polymers within the structure, or as an entity-level id (i.e. 5Y36_1) to retrieve the specific entity of interest.

**Module III** directly integrates user-provided protein sequences and associated metadata. This option enables, for example, integration of user-generated metagenome-derived proteins as well as specific curated resource datasets. For example, third-party datasets packaged with maniFasta for optional inclusion by users, include Obelisk protein sequences^36^, dietary dairy protein sequences^37,38^, and *Trichomonas tenax* protein sequences^39^, some of which are included in the AllOralsDB and described further below. The user directly supplies the desired protein sequences as a local FASTA file with an optional metadata file to supply the associated protein TaxID, genus and species for the protein sequence. The first column of the metadata file must match the first token of the FASTA protein header. If a metadata file is not supplied, default values for the TaxID is 32644 (unknown) and the genus and species are labeled NA/NA. The full functionality allows the user to incorporate self-supplied proteins alongside the specified proteins fetched.

**Module IV** allows users to fetch and select proteins from various currently supported sources and repositories, including the following:

### Human protein sequences

Users can choose which UniProtKB collection to download, including canonical sequences, canonical sequences plus isoforms, canonical plus TrEMBL, or the total canonical isoforms and TrEMBL set. The canonical protein set is exclusively Swiss-Prot which are reviewed and manually curated, the isoforms build upon the single canonical sequence by including alternate proteins generated by alternate splicing, promoter usage, translation initiation and/or ribosomal frameshifting^40^. The TrEMBL proteins are unreviewed sequences and annotations that have been computationally processed but have not been manually reviewed. If the user selects to include human proteins but does not specify which selection then the pipeline defaults to sourcing only the canonical human proteins.

### Microbial proteins

Users can choose to include proteins encoded by bacteria and archaea represented in the *e*HOMD. We use the PROKKA set available through *e*HOMD, this dataset provides open reading frame predictions and protein product annotations for all genomes, including many for which these are lacking in NCBI (including, for example, many metagenome assembled genomes - MAGs). By downloading associated available *e*HOMD metadata tables, it is possible to link the GCA identifier (present in the protein headers and a GCA_ID_info file) to their associated Human Microbial Taxon (HMT) identifiers (present in the GCA_ID_info and homd_taxonomy files). The homd_taxonomy table links each HMT to its NCBI TaxID, as well as to curated information on “Body Site(s)” associations, which is also visually represented on the daughter pages of the *e*HOMD ecology subtree (https://homd.org/taxa/ecology_home). In addition to cross-linking this information, the maniFasta pipeline allows users to filter the *e*HOMD dataset on the basis of annotated body site associations. This function adds value for users interested in primarily oral-associated taxa, as the *e*HOMD also includes genomes from other sites (e.g. vaginal, gut, environmental) and thus the full dataset is currently nearly 20M proteins. The *e*HOMD proteins can also be further deduplicated by taxonomic level, providing a single representative genome retained at the specified taxonomic rank (family, genus or species) (see Methods, Table 1). The single representative genome is selected based on the assembly contiguity, using the genome with the lowest number of contigs to represent that taxon. In the event that multiple potential representative genomes have the same number of contigs the filter selects the genome with the largest assembly size. In this way, the user is able to limit taxonomic redundancy while maintaining breadth of coverage.

### Contaminants

The identification of common sample contaminants is supported through the built-in fetch of the common Repository of Adventitious Proteins (cRAP) dataset collection hosted on Zenodo^41^. The cRAP dataset used is defined by the user in the source-plan parser, selecting either the Cambridge Centre for Proteomics (ccp), the Global Proteome Machine (gpm), the MaxQuant-distributed list (maxquant), or a combined FASTA of all three (all)^41^. The pipeline fetches the specified pre-built FASTA file(s) and then reformats the header to the maniFasta header style and incorporates the protein sequences into the final maniFasta database FASTA and records the contaminant entry into the metadata file. The contaminants are included in the maniFasta FASTA file independent of their potential inclusion in other datasets, allowing for proteins to be identified for both their biological origin and as a potential contaminant. We note that where users include maniFasta contaminant options in their builds we suggest they disable built-in contaminant sets in downstream tools such as MaxQuant^42^.

After the user has specified all desired protein sources, the pipeline launches into phase 2, where it resolves each by generating a source plan allowing all sources to be processed through various branches and then compiled into a single internally consistent protein FASTA database and metadata table. Each source has varying levels and types of information, requiring each branch to be designed to specifically shape it into the format the single build script can synthesize into the final output. A sequential internal identifier (i.e. maniFastaDB-00000000001) is assigned to each protein sequence across all collected sources and linking the same protein across the generated FASTA and metadata table. The internal ID is the sole leading token within the FASTA, with other metadata following a space-delimited format and utilizing UniProt-style formatting and key=value pairs (i.e. OS= organism OX= TaxID, and GN= gene name), allowing for easy parsing and widespread tool use.

**Table 2.** Overview of eHOMD bacterial and archaeal dataset flavors available within maniFasta. Metadata annotations provided on *e*HOMD (https://homd.org/taxa/taxon_table) can be used to filter to taxa associated with sites of interest; here we highlight configurations representing genomes of taxa with “oral” association under “Body Site(s)”, as used in performance comparisons.

| eHOMD BUILD FLAVOR | PROTEINS | GENOMES | SPECIES | GENERA | FAMILIES | INCLUDES PROTEINS FROM |
| --- | --- | --- | --- | --- | --- | --- |
| eHOMD - complete | 18,701,740 | 8,177 | . | . | . | All eHOMD genomes. |
| eHOMD - A - all oral genomes | 10,002,019 | 4,958 | 879 | 258 | 118 | All oral eHOMD genomes. |
| eHOMD - S - oral species reps | 729,243 | 345 | 345 | 97 | 55 | A single representative genome of each oral eHOMD species. |
| eHOMD - G - oral genus reps | 204,283 | 97 | 97 | 97 | 55 | A single representative genome of each oral eHOMD genus. |
| eHOMD - F - oral family reps | 117,735 | 55 | 55 | 55 | 55 | A single representative genome of each oral eHOMD family. |

In **Phase 2** maniFasta produces two key output files - the metadata table providing all information for all proteins (the manifest) and the protein reference database (the FASTA). All protein sequences are incorporated into the database with a standardized header sequence format that is prefaced with an internally consistent ID. The header sequence also incorporates taxonomy and product information, enabling smooth parsing by downstream tools such as MaxQuant^42^. The generation of the database at this stage is directed by the configuration values either specified by the user or the pipeline defaults, which are performed by the build_db.py script. For a graphical output, the pipeline generates a sunburst graph for the various taxonomies present in the database using the program plot_taxonomy.py, rendering SVG and interactive HTML files that visualize the diversity of taxa present in the user-designed database.

Finally, in **Phase 3**, all files are packaged into a final output directory labeled with a time stamp and the user-defined run label that captures not only the manifest and the FASTA, but all necessary information to document and reproduce the run. We note that this pipeline does not include a de-duplication step, thus all input proteins are present in raw maniFasta output datasets. This approach preserves all specified input data linking each protein identifier to its source taxon and is compatible with a variety of user-preferred downstream steps including: directly as the sole database, as the primary database in cascade and two-step^5,43,44^, spectral clustering^45^, or probabilistic optimization^16^, style searches; as well as for other generalized protein sequence clustering approaches such as with MMseqs2^46^.

We highlight that an important design feature of maniFasta is the collection of prepackaged inputs that are “ready to go” with the tool on the GitHub repository, and to which users and the scientific community can contribute. These inputs represent curated proteome and protein accession lists, and paired FASTA and metadata TSV datasets. In our characterization of the AllOralsDB below we describe the integration of initial prepackaged inputs, highlighting the utility of this function for collecting unique datasets that may otherwise be missed because they are distributed across various websites, public data deposits (such as Zenodo), or only available through supplementary data files associated with manuscripts.

### AllOralsDB - a phylogenetically comprehensive reference database for oral metaproteomics

Saliva is an accessible biomaterial that offers opportunities for insights into health. Application of metaproteomics to salivary diagnostics, oral microbial ecology, and oral paleoproteomics^47^, has revealed aspects of ancient human health and diets^37,38,48–50^ and modern associations between microbes and disease. The reference databases constructed for salivary and oral metaproteomics most commonly include human proteins (from the UniProtKB human proteome) and bacterial proteins (from the *e*HOMD^28,51–55^). Yet, the oral microbiome also includes diverse additional taxa not represented by these datasets, and the growing size and mixed composition of the *e*HOMD (now thousands of bacterial genomes) inflate database size and dilute ecosystem relevance. The AllOralsDB offers a solution in representing the breadth of taxa found in the oral microbiome, while also taking care to limit overall database size (Table 3). We highlight included taxa below and refer the reader to https://www.homd.org/ftp/AllOralsDB/ to download the full database and build details.

**Table 3.**
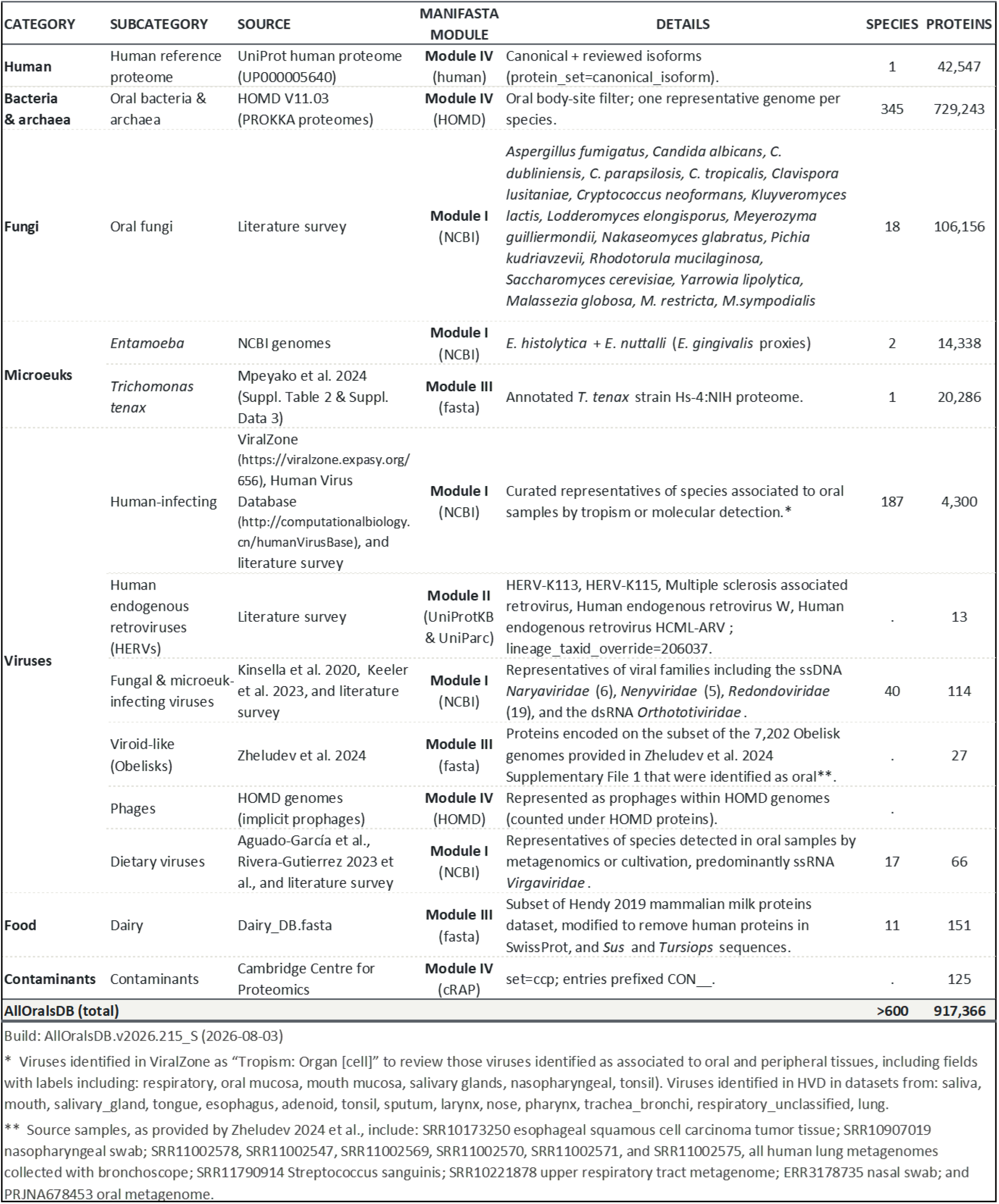
Composition and construction of the AllOralsDB (flavor S). Build: AllOralsDB.v2026.215_S.

### Human proteins

The AllOralsDB includes the UniProtKB canonical reviewed (Swiss-Prot) human proteome, as well as isoform sequences.

### Bacteria and archaea

The AllOralsDB is populated with bacterial and archaeal proteins using data available in the *e*HOMD, filtered to select only: 1) genomes annotated with “oral” “Body Site(s)” annotations in the *e*HOMD Taxon Tables, and 2) a single representative genome for each species.

### Fungi

The AllOralsDB includes fungal species previously cultured or otherwise reported in the oral mycobiome literature^56–60^. The most prevalent fungal species in the mouth is *Candida albicans*, other fungi can also be detected in abundance in some individuals, in association with oral cancers and other disease states.

### Microeukaryotes

The AllOralsDB includes sequences representing two genera of anaerobic microeukaryotes adapted to thrive in the low oxygen conditions of the human oral subgingival crevice, *Trichomonas* and *Entamoeba. Trichomonas tenax* proteins included in the database are those published by Mpeyako et al.^39^. No genome is available for *Entamoeba gingivalis*, an organism suggested to be associated with periodontal disease and the presumed host^61^ of a virus recently implicated as contributing to Sjögren’s disease^62^. The AllOralsDB therefore includes as proxies^63^ the proteomes of sequenced relatives, *E. histolytica* and *E. nuttalli* (the latter reported as detected in salivary sample metagenomes^64^), until the *E. gingivalis* genome becomes available.

### Viruses & related elements

The AllOralsDB includes sequences of phylogenetically diverse viruses and virus-like entities infecting microbial taxa of the oral microbiome and humans. Saliva contains 10^8^ virus-like particles per milliliter^65^ and salivary metaproteomics holds potential to detect human-infecting viruses^44^ as well as those infecting oral microbes.

#### Viruses, viral-satellites, and retroviruses infecting humans

Hundreds of virus species infect humans, a subset of these have oral tropism or may be detectable in the oral cavity and saliva. To define the list for inclusion in the AllOralsDB we draw on literature on the human oral virome^57,66–75^, as well as on two public databases that document associations of viral taxa with specific human tissue types, Viralzone^76^ and the Human Virus Database^77^. The AllOralsDB also includes human endogenous retrovirus protein sequences (HERVs), including from two HERV-K members that are variably present across individuals^78–80^ and have been investigated in association with diseases, including Sjögren’s^81^. Although the majority of ERVs in the human genome are degraded, some have been shown to be capable of forming particles^82^, as well as of being activated during infections by other viruses^79^.

#### Viruses infecting fungi & other microeukaryotes

Several families of viruses are known to infect human-associated microeukaryotes and fungi and are represented in the AllOralsDB. Diverse circular single-stranded DNA (ssDNA) “CRESS” viruses^83^ infect *Entamoeba*, including members of the *Naryaviridae* and *Nenyaviridae*^*84*^, and *Redondoviridae*^*61*^. Notably, the *Redondoviridae* are associated with the human oro-respiratory tract, particularly in periodontitis and critical illness^85^, and the *Vientovirus* redondoviruses have been implicated as contributing to Sjögren’s disease autoimmunity through capsid protein-based mimicking of autoantigens^62^. Diverse members of the dsRNA *Orthototiviridae* infect *Trichomonas*^*86*^ and numerous fungal species including *Malassezia*^*87,88*^, *Pichia*^*89*^, *Scheffersomyces*^*90*^ and *Saccharomyces*. Evidence of totivirus-like genes integrated in *Candida* and other fungal genomes^90^, as well as continuing discovery of diverse RNA viruses and viroid-like elements in fungi^91^, suggests that the diversity of viruses infecting human-microbiome associated fungal species will also continue to grow.

#### Viroid-like entities that infect bacteria

Protein-coding viroid-like circular RNA entities are diverse and widespread globally^92^. A class of these elements, dubbed “Obelisks”, have recently been shown to infect bacteria in the human microbiome, including the most abundant genus in the oral microbiome, *Streptococcus*^*36*^. To include oral Obelisk sequences in the AllOralsDB, we predicted open reading frames for representative Obelisk nucleotide sequences provided in Zheludev et al.^36^ using pyrodigal^93,94^, yielding a total of 11,581 protein sequences. Manual curation of source sample metadata, included in the Zheludev et al.^36^ supplementary materials, was used to identify a subset of 27 Obelisk protein sequences from oral and oral-proximal sample datasets. The full dataset of Obelisk proteins is provided as a maniFasta pre-packaged input.

#### Viruses infecting bacteria and archaea

Viruses infecting bacteria (called bacteriophages, or phages) are the most abundant type of viruses in the human mouth. Current estimates of oral phage diversity comprise tens of thousands of potential species-level units, with all bacterial and archaeal species known or expected to serve as hosts. The initial version of the AllOralsDB includes phages implicitly, as integrated into the genomes of bacteria and archaea represented in the *e*HOMD.

#### Allochthonous viruses

Viruses associated with dietary sources are detected in oral metagenomes and viromes^95,96^. These viruses are predominantly ssRNA positive-strand viruses in the family *Virgaviridae*, including tobacco mosaic virus (cultivated from the saliva of smokers^97^) and numerous species infecting plants in the family Solanaceae (representing many plant species globally important in human diets).

### Allochthonous proteins

The AllOralsDB includes the Cambridge Centre for Proteomics’ cRAP proteins set^41^, as well as a curated subset of the custom dairy database^38^ developed for the metaproteomic study of ancient human dental calculus^37^; the unmodified dairy database is provided as a maniFasta pre-packaged input.

### Comparison of AllOralsDB flavors

To identify an optimal configuration of the AllOralsDB, we considered size and performance of each version using FragPipe to analyze a test sample from a previous study^28^ optimizing detection of microbial proteins in saliva (see Methods). In this single-sample comparison, we find that the S flavor, which includes proteins from a single representative genome for each oral species in the *e*HOMD, offers near-equivalent performance to the A flavor in terms of number of peptides identified (19,289 vs 19,809) and protein groups detected (6,059 vs 6,235), while representing only 9% of the total database size and requiring 33% of the runtime (Table 4). We therefore recommend the AllOralsDB_S flavor as the reference database for analyses of salivary metaproteomes.

**Table 4.** Comparison of AllOralsDB flavor sizes and performance, using FragPipe to evaluate a publicly available oral metaproteome sample enriched in microbial proteins (see Methods).

| AllOralsDB BUILD FLAVOR | PROTEINS (FASTA size) | # of PEPTIDES identified | # of PROTEIN GROUPS | RUNTIME (mins) |
| --- | --- | --- | --- | --- |
| AllOralsDB_F | 305,858 (197 MB) | 10,429 | 3,009 | 47.9 |
| AllOralsDB_G | 392,406 (245 MB) | 12,472 | 3,838 | 51 |
| AllOralsDB_S<br>(as in Table 3) | 917,366 (538 MB) | 19,289 | 6,059 | 55 |
| AllOralsDB_A | 10,190,142 (5,647 MB) | 19,809 | 6,235 | 168.2 |

## DISCUSSION

Metaproteomic studies can offer insights into ecosystem function, yet biological samples are often highly complex, containing proteins from organisms around the tree of life, plus their viruses. “Goldilocks” reference databases that have just the right amount of sample-relevant diversity to capture signal, but not more, offer advantages for accuracy and computational efficiency^3,6^. However, design of tailored databases also introduces challenges around reproducibility and documentation.

The work presented here was motivated by the goal of better understanding host-microbe interactions in the oral microbiome through metaproteomics, a joint effort by the HOMD and HSP teams. The evident need for tools supporting flexible database construction inspired our development of maniFasta and enabled the creation of the AllOralsDB, an evolving resource tailored for phylogenetically comprehensive oral metaproteomics. To close, we address anticipated evolution of the AllOralsDB resource and highlight key features of the maniFasta toolkit.

The AllOralsDB is both a foundational oral microbiome reference database, and a substrate for iterative empirically-guided refinement. We anticipate growth of the database as genomes for relevant organisms become available (e.g. *E. gingivalis*), through investigation of unmapped *de novo*-predicted peptides and integration of their source taxon proteins into the database, and through expansion to systematically represent oral microbial species pangenomes (vs species representatives). We also anticipate contractions of the database through empirical winnowing of rarely detected taxa and proteins. The *e*HOMD has served as a key resource for salivary metaproteomics and continues to evolve, with anticipated developments including incorporation of new taxa and enriched protein annotation pipelines; the AllOralsDB will reflect these updates as they are released.

Finally, we highlight three main ways in which maniFasta supports community-driven development of biome-specific metaproteomic reference databases as a centralizing resource and open source tool. First, as the maniFasta source_registry provides an overview of all components used to build a given database, this document provides a substrate for community-driven evolution of tailored reference sets. Second, as the maniFasta code is modularized and designed to be adaptable in integrating sourcing from new data types and structures, it can continue to expand to allow users to pull from datasets and repositories not currently represented. Third, by collecting and documenting biome-specific datasets that are dispersed across numerous resources (e.g. curated public repositories, publication supplementary files, and open-access digital repositories), maniFasta centralizes information that may otherwise be dispersed and thus lost or easily missed.

## CONCLUSIONS

It is a time of plenty for ‘omics-informed biology, with an abundance of valuable data available in the public domain, and numerous initiatives to curate and enrich these datasets through various portals and repositories. In the context of metaproteomics, recent reviews and perspectives have highlighted the need for standardization in the construction and reporting of reference databases^3,26^. We address this need by developing maniFasta, an accessible and open source tool enabling users to readily construct and document databases from heterogeneous source types. Our construction of the AllOralsDB showcases maniFasta’s utility and provides a taxonomically comprehensive reference database for the oral and salivary metaproteomics community.

## AUTHOR CONTRIBUTIONS

CH - designed and developed the tool and contributed to conception and design of the study and writing of the manuscript

AKM - contributed to development of the database and writing of the manuscript

YY and MF - contributed to testing of the tool and database and reviewed and provided feedback on the manuscript

FD, TC, and JMW - reviewed and provided feedback on the manuscript

KMK - contributed to conception and design of the study, development of the tool and the database, and writing of the manuscript

## DATA AVAILABILITY

The maniFasta code underlying this work is available through GitHub (https://github.com/KauffmanLab/maniFasta) and can also be run through the Google Colab notebook linked from the GitHub repo. The current version of the AllOralsDB, and versions used in comparisons described in the manuscript, are available for download from the *e*HOMD (https://www.homd.org/ftp/AllOralsDB/).

## ACKNOWLEDGEMENTS

We thank Jasmin Pyo for assistance with beta testing. Compute resources were provided by the Center for Computational Research at the University at Buffalo.

## REFERENCES

(1) Rodríguez-Valera, F. Environmental Genomics, the Big Picture? FEMS Microbiol. Lett. 2004, 231 (2), 153–158. 10.1016/s0378-1097(04)00006-0.

(2) Wilmes, P.; Bond, P. L. The Application of Two-Dimensional Polyacrylamide Gel Electrophoresis and Downstream Analyses to a Mixed Community of Prokaryotic Microorganisms. Environ. Microbiol. 2004, 6 (9), 911–920. 10.1111/j.1462-2920.2004.00687.x.

(3) Arikan, M.; Atabay, B. Construction of Protein Sequence Databases for Metaproteomics: A Review of the Current Tools and Databases. J. Proteome Res. 2024, 23 (12), 5250–5262. 10.1021/acs.jproteome.4c00665.

(4) Van Den Bossche, T.; Armengaud, J.; Benndorf, D.; Blakeley-Ruiz, J. A.; Brauer, M.; Cheng, K.; Creskey, M.; Figeys, D.; Grenga, L.; Griffin, T. J.; Henry, C.; Hettich, R. L.; Holstein, T.; Jagtap, P. D.; Jehmlich, N.; Kim, J.; Kleiner, M.; Kunath, B. J.; Malliet, X.; Martens, L.; Mehta, S.; Mesuere, B.; Ning, Z.; Tanca, A.; Uzzau, S.; Verschaffelt, P.; Wang, J.; Wilmes, P.; Zhang, X.; Zhang, X.; Li, L.; |Metaproteomics Initiative. The Microbiologist’s Guide to Metaproteomics. Imeta 2025, 4 (3), e70031. 10.1002/imt2.70031.

(5) Jouffret, V.; Miotello, G.; Culotta, K.; Ayrault, S.; Pible, O.; Armengaud, J. Increasing the Power of Interpretation for Soil Metaproteomics Data. Microbiome 2021, 9 (1), 195. 10.1186/s40168-021-01139-1.

(6) Lee, E. M.; Srinivasan, S.; Purvine, S. O.; Fiedler, T. L.; Leiser, O. P.; Proll, S. C.; Minot, S. S.; Deatherage Kaiser, B. L.; Fredricks, D. N. Optimizing Metaproteomics Database Construction: Lessons from a Study of the Vaginal Microbiome. mSystems 2023, 8 (4), e0067822. 10.1128/msystems.00678-22.

(7) Knudsen, G. M.; Chalkley, R. J. The Effect of Using an Inappropriate Protein Database for Proteomic Data Analysis. PLoS One 2011, 6 (6), e20873. 10.1371/journal.pone.0020873.

(8) Wu, E.; Mallawaarachchi, V.; Zhao, J.; Yang, Y.; Liu, H.; Wang, X.; Shen, C.; Lin, Y.; Qiao, L. Contigs Directed Gene Annotation (ConDiGA) for Accurate Protein Sequence Database Construction in Metaproteomics. Microbiome 2024, 12 (1), 58. 10.1186/s40168-024-01775-3.

(9) Arikan, M.; Muth, T. gNOMO2: A Comprehensive and Modular Pipeline for Integrated Multi-Omics Analyses of Microbiomes. Gigascience 2024, 13 (giae038), giae038. 10.1093/gigascience/giae038.

(10) Huang, W.; Kane, M. A. MAPLE: A Microbiome Analysis Pipeline Enabling Optimal Peptide Search and Comparative Taxonomic and Functional Analysis. J. Proteome Res. 2021, 20 (5), 2882–2894. 10.1021/acs.jproteome.1c00114.

(11) Werner, J.; Géron, A.; Kerssemakers, J.; Matallana-Surget, S. mPies: A Novel Metaproteomics Tool for the Creation of Relevant Protein Databases and Automatized Protein Annotation. Biol. Direct 2019, 14 (1), 21. 10.1186/s13062-019-0253-x.

(12) Mooradian, A. D.; van der Post, S.; Naegle, K. M.; Held, J. M. ProteoClade: A Taxonomic Toolkit for Multi-Species and Metaproteomic Analysis. PLoS Comput. Biol. 2020, 16 (3), e1007741. 10.1371/journal.pcbi.1007741.

(13) Mesuere, B.; Debyser, G.; Aerts, M.; Devreese, B.; Vandamme, P.; Dawyndt, P. The Unipept Metaproteomics Analysis Pipeline. Proteomics 2015, 15 (8), 1437–1442. 10.1002/pmic.201400361.

(14) Verschaffelt, P.; Tanca, A.; Abbondio, M.; Van Den Bossche, T.; Moortele, T. V.; Dawyndt, P.; Martens, L.; Mesuere, B. Unipept Desktop 2.0: Construction of Targeted Reference Protein Databases for Metaproteogenomics Analyses. J. Proteome Res. 2023, 22 (8), 2620–2628. 10.1021/acs.jproteome.3c00091.

(15) Arikan, M. MetaproDB (https://github.com/arikanlab/MetaproDB); xGithub.

(16) Potgieter, M. G.; Nel, A. J. M.; Fortuin, S.; Garnett, S.; Wendoh, J. M.; Tabb, D. L.; Mulder, N. J.; Blackburn, J. M. MetaNovo: An Open-Source Pipeline for Probabilistic Peptide Discovery in Complex Metaproteomic Datasets. PLoS Comput. Biol. 2023, 19 (6), e1011163. 10.1371/journal.pcbi.1011163.

(17) Goldfarb, T.; Kodali, V. K.; Pujar, S.; Brover, V.; Robbertse, B.; Farrell, C. M.; Oh, D.-H.; Astashyn, A.; Ermolaeva, O.; Haddad, D.; Hlavina, W.; Hoffman, J.; Jackson, J. D.; Joardar, V. S.; Kristensen, D.; Masterson, P.; McGarvey, K. M.; McVeigh, R.; Mozes, E.; Murphy, M. R.; Schafer, S. S.; Souvorov, A.; Spurrier, B.; Strope, P. K.; Sun, H.; Vatsan, A. R.; Wallin, C.; Webb, D.; Brister, J. R.; Hatcher, E.; Kimchi, A.; Klimke, W.; Marchler-Bauer, A.; Pruitt, K. D.; Thibaud-Nissen, F.; Murphy, T. D. NCBI RefSeq: Reference Sequence Standards through 25 Years of Curation and Annotation. Nucleic Acids Res. 2025, 53 (D1), D243–D257. 10.1093/nar/gkae1038.

(18) Fullam, A.; Letunic, I.; Schmidt, T. S. B.; Ducarmon, Q. R.; Karcher, N.; Khedkar, S.; Kuhn, M.; Larralde, M.; Maistrenko, O. M.; Malfertheiner, L.; Milanese, A.; Rodrigues, J. F. M.; Sanchis-López, C.; Schudoma, C.; Szklarczyk, D.; Sunagawa, S.; Zeller, G.; Huerta-Cepas, J.; von Mering, C.; Bork, P.; Mende, D. R. proGenomes3: Approaching One Million Accurately and Consistently Annotated High-Quality Prokaryotic Genomes. Nucleic Acids Res. 2023, 51 (D1), D760–D766. 10.1093/nar/gkac1078.

(19) Richardson, L.; Allen, B.; Baldi, G.; Beracochea, M.; Bileschi, M. L.; Burdett, T.; Burgin, J.; Caballero-Pérez, J.; Cochrane, G.; Colwell, L. J.; Curtis, T.; Escobar-Zepeda, A.; Gurbich, T. A.; Kale, V.; Korobeynikov, A.; Raj, S.; Rogers, A. B.; Sakharova, E.; Sanchez, S.; Wilkinson, D. J.; Finn, R. D. MGnify: The Microbiome Sequence Data Analysis Resource in 2023. Nucleic Acids Res. 2023, 51 (D1), D753–D759. 10.1093/nar/gkac1080.

(20) Escapa, I. F.; Chen, T.; Huang, Y.; Gajare, P.; Dewhirst, F. E.; Lemon, K. P. New Insights into Human Nostril Microbiome from the Expanded Human Oral Microbiome Database (eHOMD): A Resource for the Microbiome of the Human Aerodigestive Tract. mSystems 2018, 3 (6), e00187–18. 10.1128/mSystems.00187-18.

(21) Joseph, S.; Aduse-Opoku, J.; Hashim, A.; Hanski, E.; Streich, R.; Knowles, S. C. L.; Pedersen, A. B.; Wade, W. G.; Curtis, M. A. A 16S rRNA Gene and Draft Genome Database for the Murine Oral Bacterial Community. mSystems 2021, 6 (1), e01222–20. 10.1128/mSystems.01222-20.

(22) Sunagawa, S.; Coelho, L. P.; Chaffron, S.; Kultima, J. R.; Labadie, K.; Salazar, G.; Djahanschiri, B.; Zeller, G.; Mende, D. R.; Alberti, A.; Cornejo-Castillo, F. M.; Costea, P. I.; Cruaud, C.; d’Ovidio, F.; Engelen, S.; Ferrera, I.; Gasol, J. M.; Guidi, L.; Hildebrand, F.; Kokoszka, F.; Lepoivre, C.; Lima-Mendez, G.; Poulain, J.; Poulos, B. T.; Royo-Llonch, M.; Sarmento, H.; Vieira-Silva, S.; Dimier, C.; Picheral, M.; Searson, S.; Kandels-Lewis, S.; Tara Oceans coordinators; Bowler, C.; de Vargas, C.; Gorsky, G.; Grimsley, N.; Hingamp, P.; Iudicone, D.; Jaillon, O.; Not, F.; Ogata, H.; Pesant, S.; Speich, S.; Stemmann, L.; Sullivan, M. B.; Weissenbach, J.; Wincker, P.; Karsenti, E.; Raes, J.; Acinas, S. G.; Bork, P. Ocean Plankton. Structure and Function of the Global Ocean Microbiome. Science 2015, 348 (6237), 1261359. 10.1126/science.1261359.

(23) Almeida, A.; Nayfach, S.; Boland, M.; Strozzi, F.; Beracochea, M.; Shi, Z. J.; Pollard, K. S.; Sakharova, E.; Parks, D. H.; Hugenholtz, P.; Segata, N.; Kyrpides, N. C.; Finn, R. D. A Unified Catalog of 204,938 Reference Genomes from the Human Gut Microbiome. Nat. Biotechnol. 2021, 39 (1), 105–114. 10.1038/s41587-020-0603-3.

(24) Sun, Z.; Ning, Z.; Cheng, K.; Duan, H.; Wu, Q.; Mayne, J.; Figeys, D. MetaPep: A Core Peptide Database for Faster Human Gut Metaproteomics Database Searches. Comput. Struct. Biotechnol. J. 2023, 21, 4228–4237. 10.1016/j.csbj.2023.08.025.

(25) Rigden, D. J.; Fernández, X. M. The 2026 Nucleic Acids Research Database Issue and the Online Molecular Biology Database Collection. Nucleic Acids Res. 2026, 54 (D1), D1–D9. 10.1093/nar/gkaf1427.

(26) Van Den Bossche, T.; Wolf, M.; Armengaud, J.; Arntzen, M. Ø.; Benndorf, D.; Figeys, D.; Grenga, L.; Hettich, R. L.; John, N. S.; Jagtap, P.; Jehmlich, N.; Kleiner, M.; Kunath, B. J.; Li, L.; Lipton, M.; Mesuere, B.; Neely, B. A.; Ning, Z.; Pedersen, A. L.; Perez-Riverol, Y.; Rajan, J.; Schallert, K.; Seifert, J.; Uzzau, S.; Verschaffelt, P.; Wilmes, P.; Vizcaíno, J. A.; Martens, L.; Heyer, R. The Need for Standardization and Improved Open (meta)data Practices in Metaproteomics. Microbiome 2026, 14 (1), 185. 10.1186/s40168-026-02455-0.

(27) Wilkinson, M. D.; Dumontier, M.; Aalbersberg, I. J. J.; Appleton, G.; Axton, M.; Baak, A.; Blomberg, N.; Boiten, J.-W.; da Silva Santos, L. B.; Bourne, P. E.; Bouwman, J.; Brookes, A. J.; Clark, T.; Crosas, M.; Dillo, I.; Dumon, O.; Edmunds, S.; Evelo, C. T.; Finkers, R.; Gonzalez-Beltran, A.; Gray, A. J. G.; Groth, P.; Goble, C.; Grethe, J. S.; Heringa, J.; ‘t Hoen, P. A. C.; Hooft, R.; Kuhn, T.; Kok, R.; Kok, J.; Lusher, S. J.; Martone, M. E.; Mons, A.; Packer, A. L.; Persson, B.; Rocca-Serra, P.; Roos, M.; van Schaik, R.; Sansone, S.-A.; Schultes, E.; Sengstag, T.; Slater, T.; Strawn, G.; Swertz, M. A.; Thompson, M.; van der Lei, J.; van Mulligen, E.; Velterop, J.; Waagmeester, A.; Wittenburg, P.; Wolstencroft, K.; Zhao, J.; Mons, B. The FAIR Guiding Principles for Scientific Data Management and Stewardship. Sci. Data 2016, 3 (1), 160018. 10.1038/sdata.2016.18.

(28) Yuan, J.; Sun, B.; Li, M.; Yang, C.; Zhang, L.; Chen, N.; Chen, F.; Li, L. OSaMPle Workflow for Salivary Metaproteomics Analysis Reveals Dysbiosis in Inflammatory Bowel Disease Patients. NPJ Biofilms Microbiomes 2025, 11 (1), 63. 10.1038/s41522-025-00692-z.

(29) Nesvizhskii, A. I.; Keller, A.; Kolker, E.; Aebersold, R. A Statistical Model for Identifying Proteins by Tandem Mass Spectrometry. Anal. Chem. 2003, 75 (17), 4646–4658. 10.1021/ac0341261.

(30) da Veiga Leprevost, F.; Haynes, S. E.; Avtonomov, D. M.; Chang, H.-Y.; Shanmugam, A. K.; Mellacheruvu, D.; Kong, A. T.; Nesvizhskii, A. I. Philosopher: A Versatile Toolkit for Shotgun Proteomics Data Analysis. Nat. Methods 2020, 17 (9), 869–870. 10.1038/s41592-020-0912-y.

(31) The UniProt Consortium. UniProt: The Universal Protein Knowledgebase. Nucleic Acids Res. 2017, 45 (D1), D158–D169. 10.1093/nar/gkw1099.

(32) Sayers, E. W.; Bolton, E. E.; Fine, A. M.; Kelly, C.; Kim, S.; Landrum, M.; Lathrop, S.; Malheiro, A.; Murphy, T. D.; Phan, L.; Pujar, S.; Trawick, B. W.; Schneider, V. A.; Pruitt, K. D. Database Resources of the National Center for Biotechnology Information in 2026. Nucleic Acids Res. 2026, 54 (D1), D20–D27. 10.1093/nar/gkaf1060.

(33) Goodman, R. E.; Ebisawa, M.; Ferreira, F.; Sampson, H. A.; van Ree, R.; Vieths, S.; Baumert, J. L.; Bohle, B.; Lalithambika, S.; Wise, J.; Taylor, S. L. AllergenOnline: A Peer-Reviewed, Curated Allergen Database to Assess Novel Food Proteins for Potential Cross-Reactivity. Mol. Nutr. Food Res. 2016, 60 (5), 1183–1198. 10.1002/mnfr.201500769.

(34) Leinonen, R.; Diez, F. G.; Binns, D.; Fleischmann, W.; Lopez, R.; Apweiler, R. UniProt Archive. Bioinformatics 2004, 20 (17), 3236–3237. 10.1093/bioinformatics/bth191.

(35) Berman, H. M.; Westbrook, J.; Feng, Z.; Gilliland, G.; Bhat, T. N.; Weissig, H.; Shindyalov, I. N.; Bourne, P. E. The Protein Data Bank. Nucleic Acids Res. 2000, 28 (1), 235–242. 10.1093/nar/28.1.235.

(36) Zheludev, I. N.; Edgar, R. C.; Lopez-Galiano, M. J.; de la Peña, M.; Babaian, A.; Bhatt, A. S.; Fire, A. Z. Viroid-like Colonists of Human Microbiomes. Cell 2024, 187 (23), 6521–6536.e18. 10.1016/j.cell.2024.09.033.

(37) Wilkin, S.; Ventresca Miller, A.; Taylor, W. T. T.; Miller, B. K.; Hagan, R. W.; Bleasdale, M.; Scott, A.; Gankhuyg, S.; Ramsøe, A.; Uliziibayar, S.; Trachsel, C.; Nanni, P.; Grossmann, J.; Orlando, L.; Horton, M.; Stockhammer, P. W.; Myagmar, E.; Boivin, N.; Warinner, C.; Hendy, J. Dairy Pastoralism Sustained Eastern Eurasian Steppe Populations for 5,000 Years. Nat. Ecol. Evol. 2020, 4 (3), 346–355. 10.1038/s41559-020-1120-y.

(38) Hendy, J. R. (creator). Fasta File of Custom Dairy Database Associated with Publication “Dairy Pastoralism Sustained Eastern Eurasian Steppe Populations for 5000 Years”. University of York. Dairy_DB(fasta). York Research Database, 2019. 10.15124/589742eb-287a-4576-a00a-30df33d9f52c.

(39) Mpeyako, L. A.; Hart, A. J.; Bailey, N. P.; Carlton, J. M.; Henrissat, B.; Sullivan, S. A.; Hirt, R. P. Comparative Genomics between Trichomonas Tenax and Trichomonas Vaginalis: CAZymes and Candidate Virulence Factors. Front. Microbiol. 2024, 15, 1437572. 10.3389/fmicb.2024.1437572.

(40) UniProt Consortium. UniProt: The Universal Protein Knowledgebase in 2025. Nucleic Acids Res. 2025, 53 (D1), D609–D617. 10.1093/nar/gkae1010.

(41) Gatto, L. Proteomics Contaminant Databases, 2025. 10.5281/ZENODO.15115102.

(42) Tyanova, S.; Temu, T.; Cox, J. The MaxQuant Computational Platform for Mass Spectrometry-Based Shotgun Proteomics. Nat. Protoc. 2016, 11 (12), 2301–2319. 10.1038/nprot.2016.136.

(43) Jagtap, P.; Goslinga, J.; Kooren, J. A.; McGowan, T.; Wroblewski, M. S.; Seymour, S. L.; Griffin, T. J. A Two-Step Database Search Method Improves Sensitivity in Peptide Sequence Matches for Metaproteomics and Proteogenomics Studies. Proteomics 2013, 13 (8), 1352–1357. 10.1002/pmic.201200352.

(44) Lozano, C.; Pible, O.; Eschlimann, M.; Giraud, M.; Debroas, S.; Gaillard, J.-C.; Bellanger, L.; Taysse, L.; Armengaud, J. Universal Identification of Pathogenic Viruses by Liquid Chromatography Coupled with Tandem Mass Spectrometry Proteotyping. Mol. Cell. Proteomics 2024, 23 (10), 100822. 10.1016/j.mcpro.2024.100822.

(45) Cheng, K.; Ning, Z.; Zhang, X.; Li, L.; Liao, B.; Mayne, J.; Stintzi, A.; Figeys, D. MetaLab: An Automated Pipeline for Metaproteomic Data Analysis. Microbiome 2017, 5 (1), 157. 10.1186/s40168-017-0375-2.

(46) Steinegger, M.; Söding, J. MMseqs2 Enables Sensitive Protein Sequence Searching for the Analysis of Massive Data Sets. Nat. Biotechnol. 2017, 35 (11), 1026–1028. 10.1038/nbt.3988.

(47) Warinner, C.; Korzow Richter, K.; Collins, M. J. Paleoproteomics. Chem. Rev. 2022, 122 (16), 13401–13446. 10.1021/acs.chemrev.1c00703.

(48) Bleasdale, M.; Boivin, N.; Richter, K. K. Oral Signature Screening Database for Palaeoproteomic Analyses of Dental Calculus, 2020. 10.5281/ZENODO.3698271.

(49) Bleasdale, M.; Richter, K. K.; Janzen, A.; Brown, S.; Scott, A.; Zech, J.; Wilkin, S.; Wang, K.; Schiffels, S.; Desideri, J.; Besse, M.; Reinold, J.; Saad, M.; Babiker, H.; Power, R. C.; Ndiema, E.; Ogola, C.; Manthi, F. K.; Zahir, M.; Petraglia, M.; Trachsel, C.; Nanni, P.; Grossmann, J.; Hendy, J.; Crowther, A.; Roberts, P.; Goldstein, S. T.; Boivin, N. Ancient Proteins Provide Evidence of Dairy Consumption in Eastern Africa. Nat. Commun. 2021, 12 (1), 632. 10.1038/s41467-020-20682-3.

(50) Jersie-Christensen, R. R.; Lanigan, L. T.; Lyon, D.; Mackie, M.; Belstrøm, D.; Kelstrup, C. D.; Fotakis, A. K.; Willerslev, E.; Lynnerup, N.; Jensen, L. J.; Cappellini, E.; Olsen, J. V. Quantitative Metaproteomics of Medieval Dental Calculus Reveals Individual Oral Health Status. Nat. Commun. 2018, 9 (1), 4744. 10.1038/s41467-018-07148-3.

(51) Jagtap, P.; McGowan, T.; Bandhakavi, S.; Tu, Z. J.; Seymour, S.; Griffin, T. J.; Rudney, J. D. Deep Metaproteomic Analysis of Human Salivary Supernatant. Proteomics 2012, 12 (7), 992–1001. 10.1002/pmic.201100503.

(52) Rudney, J. D.; Jagtap, P. D.; Reilly, C. S.; Chen, R.; Markowski, T. W.; Higgins, L.; Johnson, J. E.; Griffin, T. J. Protein Relative Abundance Patterns Associated with Sucrose-Induced Dysbiosis Are Conserved across Taxonomically Diverse Oral Microcosm Biofilm Models of Dental Caries. Microbiome 2015, 3 (1), 69. 10.1186/s40168-015-0136-z.

(53) Grassl, N.; Kulak, N. A.; Pichler, G.; Geyer, P. E.; Jung, J.; Schubert, S.; Sinitcyn, P.; Cox, J.; Mann, M. Ultra-Deep and Quantitative Saliva Proteome Reveals Dynamics of the Oral Microbiome. Genome Med. 2016, 8 (1), 44. 10.1186/s13073-016-0293-0.

(54) Rabe, A.; Gesell Salazar, M.; Michalik, S.; Fuchs, S.; Welk, A.; Kocher, T.; Völker, U. Metaproteomics Analysis of Microbial Diversity of Human Saliva and Tongue Dorsum in Young Healthy Individuals. J. Oral Microbiol. 2019, 11 (1), 1654786. 10.1080/20002297.2019.1654786.

(55) Rabe, A.; Gesell Salazar, M.; Michalik, S.; Kocher, T.; Below, H.; Völker, U.; Welk, A. Impact of Different Oral Treatments on the Composition of the Supragingival Plaque Microbiome. J. Oral Microbiol. 2022, 14 (1), 2138251. 10.1080/20002297.2022.2138251.

(56) Hong, B. Y.; Hoare, A.; Cardenas, A.; Dupuy, A. K.; Choquette, L.; Salner, A. L.; Schauer, P. K.; Hegde, U.; Peterson, D. E.; Dongari-Bagtzoglou, A.; Strausbaugh, L. D.; Diaz, P. I. The Salivary Mycobiome Contains 2 Ecologically Distinct Mycotypes. J. Dent. Res. 2020, 99 (6), 730–738. 10.1177/0022034520915879.

(57) Diaz, P. I. Subgingival Fungi, Archaea, and Viruses under the Omics Loupe. Periodontol. 2000 2021, 85 (1), 82–89. 10.1111/prd.12352.

(58) Nenciarini, S.; Renzi, S.; di Paola, M.; Meriggi, N.; Cavalieri, D. Ascomycetes Yeasts: The Hidden Part of Human Microbiome. WIREs Mech. Dis. 2024, 16 (3), e1641. 10.1002/wsbm.1641.

(59) Mukherjee, P. K.; Chandra, J.; Retuerto, M.; Sikaroodi, M.; Brown, R. E.; Jurevic, R.; Salata, R. A.; Lederman, M. M.; Gillevet, P. M.; Ghannoum, M. A. Oral Mycobiome Analysis of HIV-Infected Patients: Identification of Pichia as an Antagonist of Opportunistic Fungi. PLoS Pathog. 2014, 10 (3), e1003996. 10.1371/journal.ppat.1003996.

(60) He, S.; Chakraborty, R.; Ranganathan, S. Metaproteomic Analysis of an Oral Squamous Cell Carcinoma Dataset Suggests Diagnostic Potential of the Mycobiome. Int. J. Mol. Sci. 2023, 24 (2), 1050. 10.3390/ijms24021050.

(61) Keeler, E. L.; Merenstein, C.; Reddy, S.; Taylor, L. J.; Cobián-Güemes, A. G.; Zankharia, U.; Collman, R. G.; Bushman, F. D. Widespread, Human-Associated Redondoviruses Infect the Commensal Protozoan Entamoeba Gingivalis. Cell Host Microbe 2023, 31 (1), 58–68.e5. 10.1016/j.chom.2022.11.002.

(62) Zhang, X.; Li, Y.; Qin, Y.; Liao, Z.; Deng, C.; Chen, Y.; Li, Y.; Qian, H.; He, Y.; Chen, S.; Shi, G.; Liu, Y. Vientovirus Capsid Protein Mimics Autoantigens and Contributes to Autoimmunity in Sjögren’s Disease. Nat. Microbiol. 2025, 10 (10), 2591–2602. 10.1038/s41564-025-02115-3.

(63) Shah, T.; Fitzpatrick, J. A.; Orsburn, B. C. Backyard Proteomics: A Case Study with the Black Widow Spider. J. Proteome Res. 2025, 24 (9), 4838–4844. 10.1021/acs.jproteome.5c00342.

(64) Bao, K.; Afacan, B.; Grossmann, J.; Silbereisen, A.; Öztürk, V.-Ö.; Emingil, G.; Belibasakis, G. N.; Bostanci, N. Saliva versus All-Site Microbiome and Proteome Mapping in Periodontitis. J. Clin. Periodontol. 2025, 52 (11), 1540–1549. 10.1111/jcpe.70017.

(65) Pride, D. T.; Salzman, J.; Haynes, M.; Rohwer, F.; Davis-Long, C.; White, R. A., 3rd; Loomer, P.; Armitage, G. C.; Relman, D. A. Evidence of a Robust Resident Bacteriophage Population Revealed through Analysis of the Human Salivary Virome. ISME J. 2012, 6 (5), 915–926. 10.1038/ismej.2011.169.

(66) Corstjens, P. L. A. M.; Abrams, W. R.; Malamud, D. Detecting Viruses by Using Salivary Diagnostics. J. Am. Dent. Assoc. 2012, 143 (10 Suppl), 12S –18S. 10.14219/jada.archive.2012.0338.

(67) Corstjens, P. L. A. M.; Abrams, W. R.; Malamud, D. Saliva and Viral Infections. Periodontol. 2000 2016, 70 (1), 93–110. 10.1111/prd.12112.

(68) Martínez, A.; Kuraji, R.; Kapila, Y. L. The Human Oral Virome: Shedding Light on the Dark Matter. Periodontol. 2000 2021, 87 (1), 282–298. 10.1111/prd.12396.

(69) Paietta, E. N.; Kraberger, S.; Custer, J. M.; Vargas, K. L.; Espy, C.; Ehmke, E.; Yoder, A. D.; Varsani, A. Characterization of Diverse Anelloviruses, Cressdnaviruses, and Bacteriophages in the Human Oral DNA Virome from North Carolina (USA). Viruses 2023, 15 (9), 1821. 10.3390/v15091821.

(70) Hernandez-Kapila, Y. L.; Kim, R. Y.; Gokhale, S. S.; Lin, Y.-L.; Dommisch, H.; Slots, J.; Weisenberger, D. J. Oral Virome in Health and Disease. J. Am. Dent. Assoc. 2026. 10.1016/j.adaj.2026.01.019.

(71) Bottalico, D.; Chen, Z.; Dunne, A.; Ostoloza, J.; McKinney, S.; Sun, C.; Schlecht, N. F.; Fatahzadeh, M.; Herrero, R.; Schiffman, M.; Burk, R. D. The Oral Cavity Contains Abundant Known and Novel Human Papillomaviruses from the Betapapillomavirus and Gammapapillomavirus Genera. J. Infect. Dis. 2011, 204 (5), 787–792. 10.1093/infdis/jir383.

(72) Ure, A. E.; Forslund, O. Characterization of Human Papillomavirus Type 154 and Tissue Tropism of Gammapapillomaviruses. PLoS One 2014, 9 (2), e89342. 10.1371/journal.pone.0089342.

(73) Modha, S.; Hughes, J.; Orton, R. J.; Lytras, S. Expanding the Genomic Diversity of Human Anelloviruses. Virus Evol. 2025, 11 (1), veaf002. 10.1093/ve/veaf002.

(74) Weller, M. L.; Gardener, M. R.; Bogus, Z. C.; Smith, M. A.; Astorri, E.; Michael, D. G.; Michael, D. A.; Zheng, C.; Burbelo, P. D.; Lai, Z.; Wilson, P. A.; Swaim, W.; Handelman, B.; Afione, S. A.; Bombardieri, M.; Chiorini, J. A. Hepatitis Delta Virus Detected in Salivary Glands of Sjögren’s Syndrome Patients and Recapitulates a Sjögren’s Syndrome-like Phenotype in Vivo. Pathog. Immun. 2016, 1 (1), 12–40. 10.20411/pai.v1i1.72.

(75) Bergner, L. M.; Orton, R. J.; Broos, A.; Tello, C.; Becker, D. J.; Carrera, J. E.; Patel, A. H.; Biek, R.; Streicker, D. G. Diversification of Mammalian Deltaviruses by Host Shifting. Proc. Natl. Acad. Sci. U. S. A. 2021, 118 (3), e2019907118. 10.1073/pnas.2019907118.

(76) Hulo, C.; de Castro, E.; Masson, P.; Bougueleret, L.; Bairoch, A.; Xenarios, I.; Le Mercier, P. ViralZone: A Knowledge Resource to Understand Virus Diversity. Nucleic Acids Res. 2011, 39 (Database issue), D576–D582. 10.1093/nar/gkq901.

(77) Ye, S.; Lu, C.; Qiu, Y.; Zheng, H.; Ge, X.; Wu, A.; Xia, Z.; Jiang, T.; Zhu, H.; Peng, Y. An Atlas of Human Viruses Provides New Insights into Diversity and Tissue Tropism of Human Viruses. Bioinformatics 2022, 38 (11), 3087–3093. 10.1093/bioinformatics/btac275.

(78) Wildschutte, J. H.; Williams, Z. H.; Montesion, M.; Subramanian, R. P.; Kidd, J. M.; Coffin, J. M. Discovery of Unfixed Endogenous Retrovirus Insertions in Diverse Human Populations. Proc. Natl. Acad. Sci. U. S. A. 2016, 113 (16), E2326–E2334. 10.1073/pnas.1602336113.

(79) Evans, E. F.; Saraph, A.; Tokuyama, M. Transactivation of Human Endogenous Retroviruses by Viruses. Viruses 2024, 16 (11), 1649. 10.3390/v16111649.

(80) Kyriakou, E.; Magiorkinis, G. Compilation of All Known HERV-K HML-2 Proviral Integrations. Mob. DNA 2025, 16 (1), 21. 10.1186/s13100-025-00359-8.

(81) Moyes, D. L.; Martin, A.; Sawcer, S.; Temperton, N.; Worthington, J.; Griffiths, D. J.; Venables, P. J. The Distribution of the Endogenous Retroviruses HERV-K113 and HERV-K115 in Health and Disease. Genomics 2005, 86 (3), 337–341. 10.1016/j.ygeno.2005.06.004.

(82) Boller, K.; Schönfeld, K.; Lischer, S.; Fischer, N.; Hoffmann, A.; Kurth, R.; Tönjes, R. R. Human Endogenous Retrovirus HERV-K113 Is Capable of Producing Intact Viral Particles. J. Gen. Virol. 2008, 89 (Pt 2), 567–572. 10.1099/vir.0.83534-0.

(83) Krupovic, M.; Varsani, A. Naryaviridae, Nenyaviridae, and Vilyaviridae: Three New Families of Single-Stranded DNA Viruses in the Phylum Cressdnaviricota. Arch. Virol. 2022, 167 (12), 2907–2921. 10.1007/s00705-022-05557-w.

(84) Kinsella, C. M.; Bart, A.; Deijs, M.; Broekhuizen, P.; Kaczorowska, J.; Jebbink, M. F.; van Gool, T.; Cotten, M.; van der Hoek, L. Entamoeba and Giardia Parasites Implicated as Hosts of CRESS Viruses. Nat. Commun. 2020, 11 (1), 4620. 10.1038/s41467-020-18474-w.

(85) Abbas, A. A.; Taylor, L. J.; Dothard, M. I.; Leiby, J. S.; Fitzgerald, A. S.; Khatib, L. A.; Collman, R. G.; Bushman, F. D. Redondoviridae, a Family of Small, Circular DNA Viruses of the Human Oro-Respiratory Tract Associated with Periodontitis and Critical Illness. Cell Host Microbe 2019, 25 (5), 719–729.e4. 10.1016/j.chom.2019.04.001.

(86) Tai, J. H.; Ip, C. F. The cDNA Sequence of Trichomonas Vaginalis Virus-T1 Double-Stranded RNA. Virology 1995, 206 (1), 773–776. 10.1016/s0042-6822(95)80008-5.

(87) Applen Clancey, S.; Ruchti, F.; LeibundGut-Landmann, S.; Heitman, J.; Ianiri, G. A Novel Mycovirus Evokes Transcriptional Rewiring in the Fungus Malassezia and Stimulates Beta Interferon Production in Macrophages. MBio 2020, 11 (5). 10.1128/mBio.01534-20.

(88) Park, M.; Cho, Y.-J.; Kim, D.; Yang, C.-S.; Lee, S. M.; Dawson, T. L., Jr; Nakamizo, S.; Kabashima, K.; Lee, Y. W.; Jung, W. H. A Novel Virus Alters Gene Expression and Vacuolar Morphology in Malassezia Cells and Induces a TLR3-Mediated Inflammatory Immune Response. MBio 2020, 11 (5). 10.1128/mBio.01521-20.

(89) Lee, M. D.; Creagh, J. W.; Fredericks, L. R.; Crabtree, A. M.; Patel, J. S.; Rowley, P. A. The Characterization of a Novel Virus Discovered in the Yeast Pichia Membranifaciens. Viruses 2022, 14 (3), 594. 10.3390/v14030594.

(90) Taylor, D. J.; Bruenn, J. The Evolution of Novel Fungal Genes from Non-Retroviral RNA Viruses. BMC Biol. 2009, 7 (1), 88. 10.1186/1741-7007-7-88.

(91) Forgia, M.; Navarro, B.; Daghino, S.; Cervera, A.; Gisel, A.; Perotto, S.; Aghayeva, D. N.; Akinyuwa, M. F.; Gobbi, E.; Zheludev, I. N.; Edgar, R. C.; Chikhi, R.; Turina, M.; Babaian, A.; Di Serio, F.; de la Peña, M. Hybrids of RNA Viruses and Viroid-like Elements Replicate in Fungi. Nat. Commun. 2023, 14 (1), 2591. 10.1038/s41467-023-38301-2.

(92) Lee, B. D.; Neri, U.; Roux, S.; Wolf, Y. I.; Camargo, A. P.; Krupovic, M.; RNA Virus Discovery Consortium; Simmonds, P.; Kyrpides, N.; Gophna, U.; Dolja, V. V.; Koonin, E. V. Mining Metatranscriptomes Reveals a Vast World of Viroid-like Circular RNAs. Cell 2023, 186 (3), 646–661.e4. 10.1016/j.cell.2022.12.039.

(93) Hyatt, D.; Chen, G.-L.; Locascio, P. F.; Land, M. L.; Larimer, F. W.; Hauser, L. J. Prodigal: Prokaryotic Gene Recognition and Translation Initiation Site Identification. BMC Bioinformatics 2010, 11, 119. 10.1186/1471-2105-11-119.

(94) Larralde, M. Pyrodigal: Python Bindings and Interface to Prodigal, an Efficient Method for Gene Prediction in Prokaryotes. J. Open Source Softw. 2022, 7 (72), 4296. 10.21105/joss.04296.

(95) Aguado-García, Y.; Taboada, B.; Morán, P.; Rivera-Gutiérrez, X.; Serrano-Vázquez, A.; Iša, P.; Rojas-Velázquez, L.; Pérez-Juárez, H.; López, S.; Torres, J.; Ximénez, C.; Arias, C. F. Tobamoviruses Can Be Frequently Present in the Oropharynx and Gut of Infants during Their First Year of Life. Sci. Rep. 2020, 10 (1), 13595. 10.1038/s41598-020-70684-w.

(96) Rivera-Gutiérrez, X.; Morán, P.; Taboada, B.; Serrano-Vázquez, A.; Isa, P.; Rojas-Velázquez, L.; Pérez-Juárez, H.; López, S.; Torres, J.; Ximénez, C.; Arias, C. F. The Fecal and Oropharyngeal Eukaryotic Viromes of Healthy Infants during the First Year of Life Are Personal. Sci. Rep. 2023, 13 (1), 938. 10.1038/s41598-022-26707-9.

(97) Balique, F.; Colson, P.; Raoult, D. Tobacco Mosaic Virus in Cigarettes and Saliva of Smokers. J. Clin. Virol. 2012, 55 (4), 374–376. 10.1016/j.jcv.2012.08.012.

